# Bacterial sepsis triggers stronger transcriptomic immune responses in larger primates

**DOI:** 10.1101/2023.07.11.548565

**Authors:** Ryan McMinds, Rays H.Y. Jiang, Swamy R. Adapa, Emily Cornelius Ruhs, Rachel A. Munds, Jennifer W. Leiding, Cynthia J. Downs, Lynn B. Martin

## Abstract

Empirical data relating body mass to immune defense against infections remain limited. Although the Metabolic Theory of Ecology predicts that larger organisms would have weaker immune responses, recent studies have suggested that the opposite may be true. These discoveries have led to the Safety Factor hypothesis, which proposes that larger organisms have evolved stronger immune defenses because they carry greater risks of exposure to pathogens and parasites. In this study, we simulated sepsis by exposing blood from nine primate species to a bacterial lipopolysaccharide (LPS), measured the relative expression of immune and other genes using RNAseq, and fit phylogenetic models to determine how gene expression was related to body mass. In contrast to non-immune-annotated genes, we discovered hypermetric scaling in the LPS-induced expression of innate immune genes, such that large primates had a disproportionately greater increase in gene expression of immune genes compared to small primates. Hypermetric immune gene expression appears to support the Safety Factor hypothesis, though this pattern may represent a balanced evolutionary mechanism to compensate for lower per-transcript immunological effectiveness. This study contributes to the growing body of immune allometry research, highlighting its importance in understanding the complex interplay between body size and immunity over evolutionary timescales.

**Author Summary:** Understanding the relationship between an animal’s size and its ability to defend against disease can inform predictions about evolutionary tradeoffs and susceptibility to infection. Two major theories – the Metabolic Theory of Ecology (MTE) and the Safety Factor hypothesis – offer opposing views on how body size influences it immune defenses. In this study, we compared the immune gene expression of nine primate species to a simulated bacterial infection. We found that larger species mounted a stronger transcriptional immune response, consistent with either the Safety Factor hypothesis or an evolutionary pressure to compensate for and balance the effects expected in the context of the MTE.

## Introduction

The body size of organisms profoundly affects most of their life processes [1]. Partly, these effects are due to physical forces affecting how materials are moved into and out of cells, but evolutionary factors, too, can influence how size affects life [2]. For instance, reaching a large size inherently requires time, so selection should act strongly on developmental rates and age-specific breeding traits, which could lead to innovative molecular or physiological mechanisms that circumvent some physical constraints. Relationships between size and biological traits are often modeled as trait ∼ α * mass^β^, which can be linearized via log-transformation of both body mass and the response variable (i.e., log(trait) ∼ log(α) + β * log(mass)). Relationships between body mass and response variables are then represented by the slope coefficients, β, which can be estimated with linear regression. Traits such as blood volume and length of the digestive tract scale proportionally to body size (i.e., isometrically), with β approximately equal to 1 [3,4]. Other traits, such as hematocrit, also scale isometrically [5,6], but because they are measures of concentration that are inherently scaled by body mass, they have coefficients approximately equal to 0. Allometric relationships, where β deviates from the proportional expectation, are also quite common. A trait with β greater than isometry is called hyp<u>er</u>metric whereas one with β less than isometry is hyp<u>o</u>metric. A prime example of hypoallometry is organismal metabolic rate, which scales at ¾ the rate of body mass change. In other words, the total metabolic output of an organism increases as body mass increases, but at less than a one-to-one ratio. Equivalently, mass-normalized metabolic rate has β = -¼ (i.e., the metabolic rate *per unit mass* **decreases** as body mass itself increases) [7–9].

Biological scaling has been investigated extensively [6,10,11], and metabolic allometries in particular have been so powerful as to lead to the Metabolic Theory of Ecology (MTE) [7,12]. Although contentious [13–16], the MTE has been useful for everything from improving the conservation of wildlife to understanding the structure of human cities [1]. Nevertheless, the MTE so far appears to hold little utility for understanding how defenses, specifically immune system processes, relate to body size. Whereas the MTE predicts hypometric immune cell concentrations (β ≈ -¼) [17], granulocyte concentrations in birds and mammals in fact scale hypermetrically (heterophils and neutrophils scale with β = 0.19 and 0.11, respectively) [18,19]. These unexpected results suggest that other processes must come into play to explain how defenses scale with body size, and the Safety Factor hypothesis was recently offered as an eco-evolutionary explanation [20]. This hypothesis proposes that large organisms are exposed to more numerous and more diverse pathogens within and between generations, such that the risks and consequences of infection are much greater compared to smaller species. As a result, large species may be selected to evolve exceptionally strong defenses compared to small species [19].

Like most physiological processes involving energy metabolism, gene expression (both the rate of gene transcription and the abundance of gene transcripts) is predicted by the MTE to scale hypometrically. A recent study including nematodes, insects, crustaceans, and fish provides evidence that this is broadly true across the transcriptome [21], but whether immune genes are an exception to this trend is unknown. Immune gene expression is of particular interest because it could give more insight into the granulocyte hyperallometry previously described. If immune gene expression scaling is hypometric, then hypermetric granulocyte concentrations could simply act as compensation for relatively poor per-cell function. Alternatively, hypermetric immune gene expression could provide large species with disproportionately stronger defenses. Other recent studies produced data consistent with immune gene hyperallometry, but due to their limited phylogenetic sampling, they could not disambiguate allometric effects from effects of shared ancestry or other confounding variables [22,23].

In principle, the prediction for exceptionally strong defenses in large species could be fulfilled by hypermetric baseline expression (i.e., the immune system maintains *consistently* greater constitutive expression of defensive genes), by hypermetric response to infection (i.e., immune gene expression is increased disproportionately *only after detection* of a pathogen), or by both (Fig. 1A-C). In plants, large, long-lived species tend to have stronger constitutive barriers than small, short-lived species [24]. Similar patterns have also been suggested across vertebrates [25,26], and such a strategy could be necessary in the context of the slower metabolism observed in larger animals. However, greater baseline gene expression would be more costly, and the balance of these tradeoffs is unclear.

**Fig. 1.**
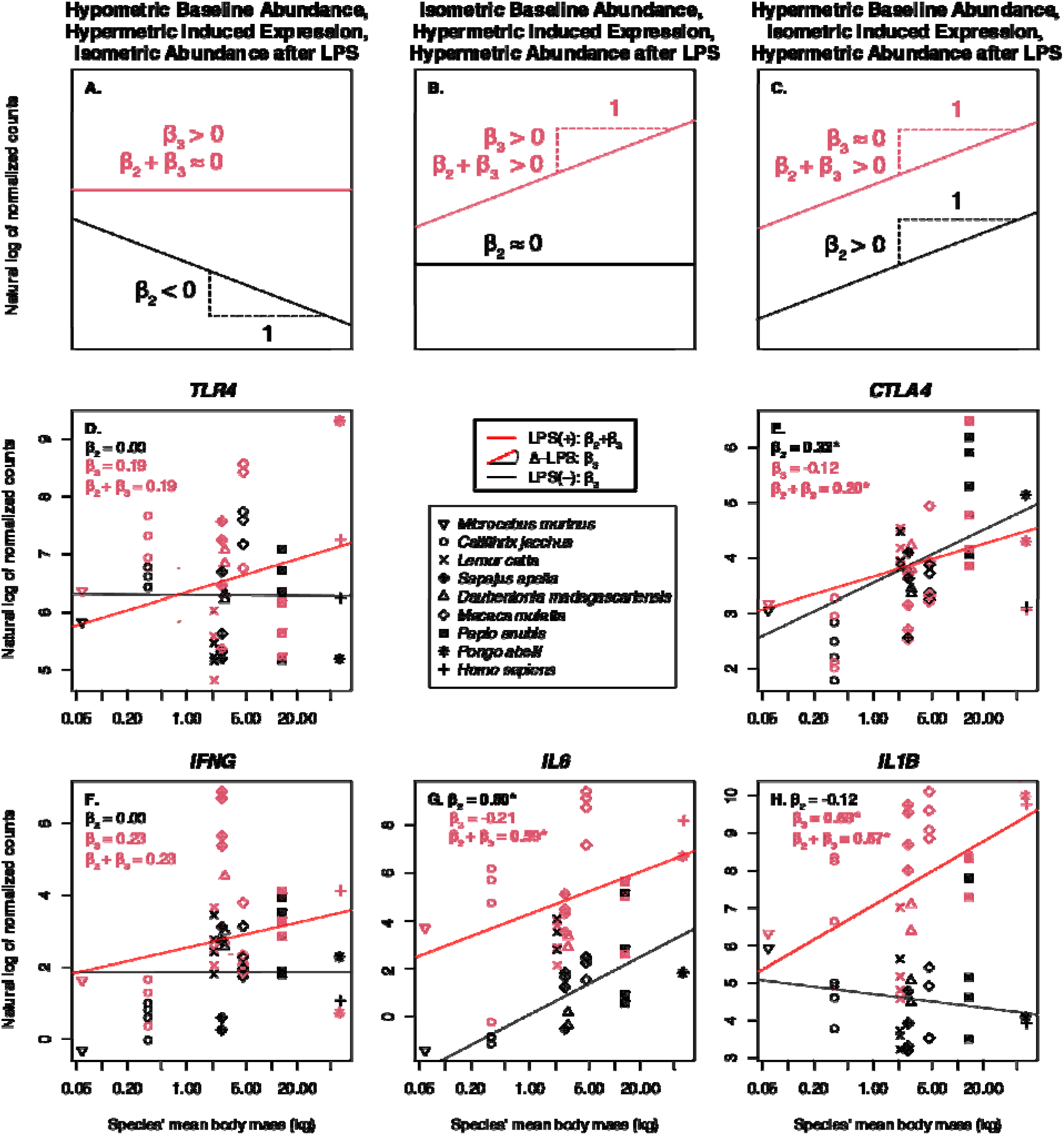
Immune genes are characterized by distinct allometries that often result in hypermetric relative transcript abundance during simulated sepsis. (**A-C**) Example expression patterns demonstrate how phylogenetic generalized linear mixed model (pGLMM) coefficients correspond to different forms of allometry. **(A)** Hypometric baseline abundance (β_2_ < 0) is compensated for with hypermetric LPS-induced expression (β_3_ > 0), resulting in isometric abundance after LPS (β_2_+ β_3_ ∼ 0). **(B)** Isometric baseline abundance (β_2_ ∼ 0) plus hypermetric LPS-induced expression (β_3_> 0) results in hypermetric abundance after LPS (β_2_ + β_3_> 0). **(C)** Hypermetric baseline abundance (β_2_ > 0) plus isometric LPS-induced expression (β_3_∼ 0) results in hypermetric abundance after LPS (β_2_ + β_3_> 0). (**D-H**) Log abundance (relative to the per-sample size factor and after adding a pseudocount of 0.5) of five example genes (HGNC gene names *TLR4*, *CTLA4*, *IFNG*, *IL6*, and *IL1B*, respectively) as a function of species’ average body mass in kilograms (plotted on log scale). Symbols correspond to species (l7= *Microcebus murinus*, ○ = *Callithrix jacchus*, × = *Lemur catta*, l7+ = *Sapajus apella*, △ = *Daubentonia madagascariensis*, l7 = *Macaca mulatta*, l7 = *Papio anubis*, * = *Pongo abelii*, + = *Homo sapiens*). Black points correspond to samples with control treatment, and red points correspond to samples with LPS treatment. Fitted lines are plotted using coefficients extracted from the pGLMMs, with dashed lines for 95% credible intervals. pGLMM coefficients are listed, with an asterisk (*) if the 95% credible interval did not include zero.

To assess the allometry of immune gene expression in these distinct contexts, we compared the transcriptomic profiles of replicate aliquots of live blood of adult male individuals from nine primate species after i) a standard dose of lipopolysaccharide (LPS) from *Escherichia coli* to simulate sepsis (LPS(+)), and ii) a null (LPS(-)) treatment of LPS-free growth media to assess baseline expression. Unlike a recent similar study [22], which included only four relatively large catarrhine species, our study included platyrrhine and strepsirrhine species as well, and spanned almost the full range of body masses of species in the Primates order (Fig. 2). We used Bayesian phylogenetic generalized linear mixed models (pGLMMs) to link raw transcript abundance data with fixed effects of the form α + β_1_ * treatment + β_2_ * log(mass) + β_3_ * treatment * log(mass), which links the allometry of baseline (LPS(-)) transcript abundance to β_2_, the allometry of LPS-induced gene expression (Δ-LPS) to β_3_, and the allometry of transcript abundance after LPS (LPS(+)) to the sum of β_2_ and β_3_ (Fig. 1). Models also included an offset to compensate for transcript compositionality, additional fixed effects to control for within-species differences in body size, and random effects to control for shared phylogenetic history, repeated measures, and statistical overdispersion. By standardizing our simulated sepsis methods and comparing transcriptomic responses in a phylogenetically informed context, we revealed that innate immune gene expression in response to simulated sepsis scales hypermetrically among Primates.

**Fig. 2.**
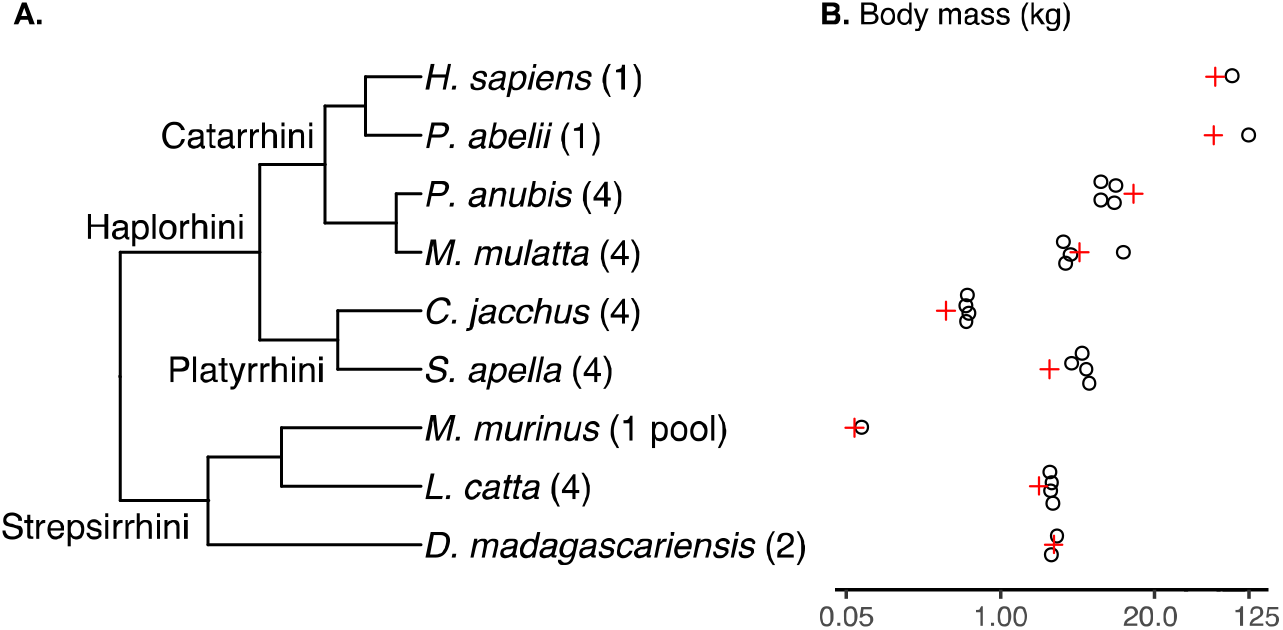
Body mass and phylogenetic relationships among samples. (**A**) Primates phylogeny derived from [27]. Tips are annotated with the numbers of individuals sampled for the present study (the pool for *M. murinus* consisted of 8 individuals). (**B**) Body mass (kg) in the context of their phylogenetic relationships (° = individuals in our study, + = species mean from [28].

## Results and Discussion

### The total immune gene transcriptional response to LPS scales hypermetrically among Primate species

To assess the overall allometric effects on immune gene expression, we first pooled all transcript counts that mapped to immune-annotated one-to-one orthologous genes. Baseline relative abundance (LPS(-)) of this sum of immune-annotated transcripts did not scale significantly differently from isometry (β_2_ ∼ -0.03), but the LPS-induced expression (Δ-LPS) did scale hypermetrically (β_3_ ∼ 0.13) with 95% credibility. Although the relative abundance of immune gene transcripts after LPS exposure (LPS(+)) did not scale with 95%-credible hyperallometry (β_2_ + β_3_ ∼ 0.10), the bulk of the posterior was positive (Table 1; Figs. 3, S1, S4-5).

**Table 1.** Pooled analyses of immune-annotated genes support hypermetric LPS-induced expression. Medians and 95% credible intervals for betas extracted from Bayesian pGLMMs fit to different pools of immune-annotated genes. 9-sp refers to analyses with all 9 sampled species, and 6-sp refers to analyses of only the 6 species with high-quality reference genomes. Orthogroups include 1:1 orthologous genes, but also genes that appear to have had duplications or deletions in the sampled species. Baseline relative abundance = LPS(-), LPS-induced expression = Δ-LPS, and relative abundance after LPS = LPS(+).

| Data subset | Gene expression context | Parameter | Posterior 2.5% CI | Posterior median | Posterior 97.5% CI |
| --- | --- | --- | --- | --- | --- |
| **9-sp, 1:1 orthologs** | LPS(-) | $\beta_2$ | -0.17 | -0.03 | 0.10 |
| **9-sp, 1:1 orthologs** | Δ-LPS | $\beta_3$ | **0.01** | 0.13 | 0.25 |
| **9-sp, 1:1 orthologs** | LPS(+) | $\beta_2 + \beta_3$ | -0.04 | 0.10 | 0.23 |
| **9-sp, orthogroups** | LPS(-) | $\beta_2$ | -0.07 | 0.05 | 0.17 |
| **9-sp, orthogroups** | Δ-LPS | $\beta_3$ | -0.0020 | 0.08 | 0.16 |
| **9-sp, orthogroups** | LPS(+) | $\beta_2 + \beta_3$ | **0.01** | 0.13 | 0.25 |
| **6-sp, 1:1 orthologs** | LPS(-) | $\beta_2$ | -0.05 | 0.02 | 0.09 |
| **6-sp, 1:1 orthologs** | Δ-LPS | $\beta_3$ | **0.04** | 0.13 | 0.23 |
| **6-sp, 1:1 orthologs** | LPS(+) | $\beta_2 + \beta_3$ | **0.08** | 0.15 | 0.22 |
| **6-sp, orthogroups** | LPS(-) | $\beta_2$ | -0.25 | -0.13 | **-0.01** |
| **6-sp, orthogroups** | Δ-LPS | $\beta_3$ | **0.01** | 0.17 | 0.34 |
| **6-sp, orthogroups** | LPS(+) | $\beta_2 + \beta_3$ | -0.07 | 0.04 | 0.16 |

**Fig. 3.**
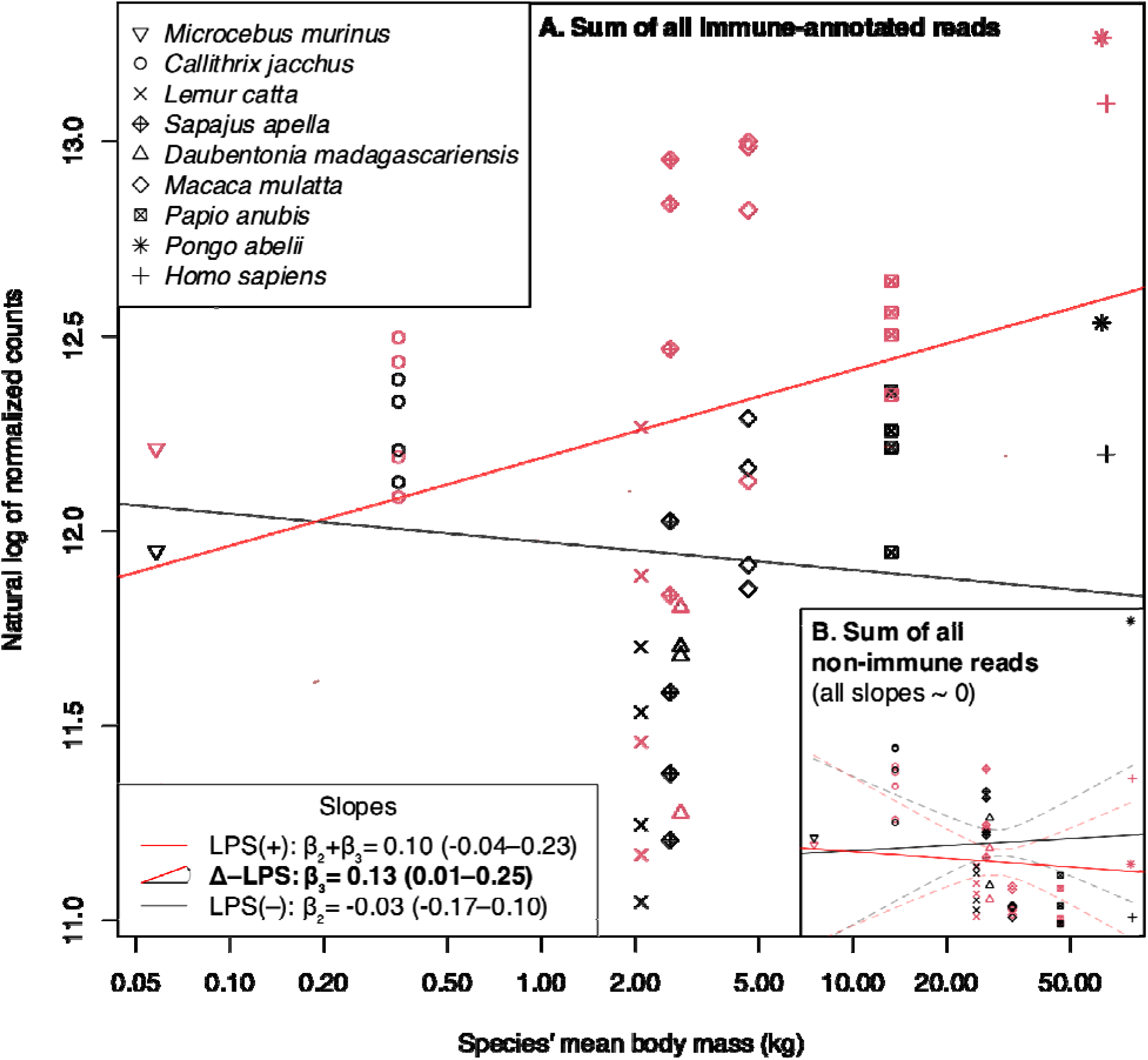
LPS-induced expression of immune genes scales hypermetrically among nine primate species. (**A**) Log abundance (relative to the per-sample size factor and after adding a pseudocount of 0.5) of all reads belonging to immune-annotated 1:1 orthologous genes as a function of species’ average body mass in kilograms (plotted on log scale). Symbols correspond to species (IZ= *Microcebus murinus*, ○ = *Callithrix jacchus*, × = *Lemur catta*, IZ+ = *Sapajus apella*, △ = *Daubentonia madagascariensis*, IZ = *Macaca mulatta*, IZ = *Papio anubis*, * = *Pongo abelii*, + = *Homo sapiens*). Black points correspond to samples with control treatment, and red points to samples with LPS treatment. Fitted lines are plotted using coefficients extracted from the pGLMMs, with dashed lines for 95% credible intervals. pGLMM coefficients are listed in the lower left legend, with the estimate followed by the 95% credible interval. (**B**) Log relative abundance of non-immune-annotated 1:1 orthologous genes.

One persistent complication inherent to comparative transcriptomics is how to deal with the evolution of gene copy number. Because gene duplication is sometimes, but not always, associated with punctuated (non-Brownian) evolution and neofunctionalization, fewer assumptions are required if we limit an analysis to only one-to-one orthologous genes, as in our primary analysis. However, varying copy number is one mechanism by which a gene may vary its expression levels, so filtering out genes with duplications and losses has the potential to bias comparative analyses of expression. We thus repeated our analysis of the overall expression of immune genes using the sum of all reads that mapped to immune orthogroups, including duplicated and paralogous genes. This analysis returned hypermetric estimates for all of baseline relative abundance (β_2_ ∼ 0.05), LPS-induced expression (β_3_∼ 0.08), and relative abundance after LPS (β_2_ + β_3_ ∼ 0.13), although only relative abundance after LPS was 95% credible (Table 1; Figs. S2-3, S6-7). Together, these analyses suggest that larger primates have disproportionally stronger responses to LPS, leading to higher relative abundances of immune gene transcripts during sepsis, than do smaller primates.

### Individual immune-annotated genes are more likely than non-immune genes to scale hypermetrically

Although the total expression of immune-annotated genes is an important measure of the total activity of cells, it is possible that such a pattern is driven by the expression of a few highly expressed genes. Thus, to assess broader trends across the transcriptome, we additionally analyzed the expression of each one-to-one orthologous gene in our dataset and classified them as hypometric, hypermetric, or of uncertain relation to body mass. Using Fisher’s exact tests on the resulting contingency tables, we found that individual immune-annotated genes were more likely than non-immune-annotated genes to scale hypermetrically in terms of their LPS-induced expression and relative abundance after LPS, but not their baseline relative abundance (Table 2; Data S2).

**Table 2.**
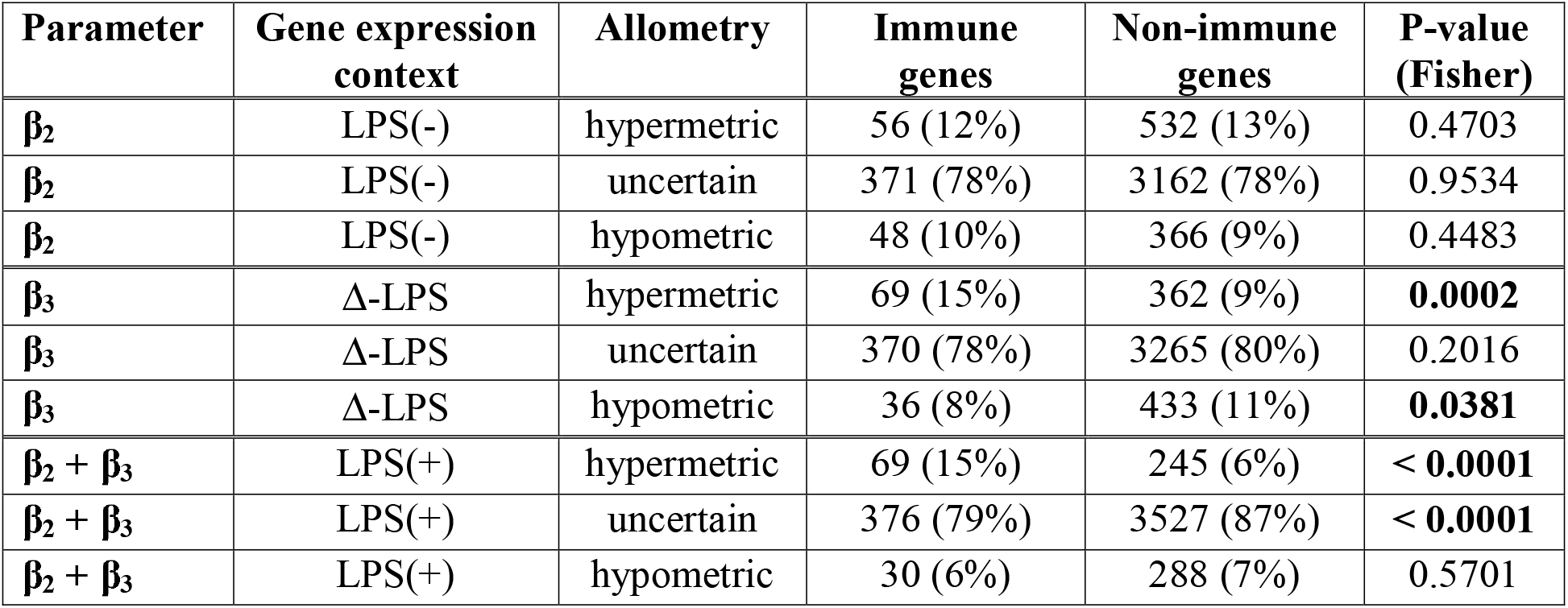
Number of immune- or non-immune-annotated genes with hypometric, hypermetric, or uncertain relation to body mass. Genes were considered hypometric in each context (baseline relative abundance = LPS(-), LPS-induced expression = Δ-LPS, and relative abundance after LPS = LPS(+)) if the pGLMM 95% credible interval for the corresponding beta was entirely negative, hypermetric if it was entirely positive, and uncertain if it included zero. P-values correspond to Fisher’s exact test comparing the proportions of immune or non-immune genes with each allometric pattern vs. the sum of the other two patterns.

| Parameter | Gene expression context | Allometry | Immune genes | Non-immune genes | P-value (Fisher) |
| --- | --- | --- | --- | --- | --- |
| $\beta_2$ | LPS(-) | hypermetric | 56 (12%) | 532 (13%) | 0.4703 |
| $\beta_2$ | LPS(-) | uncertain | 371 (78%) | 3162 (78%) | 0.9534 |
| $\beta_2$ | LPS(-) | hypometric | 48 (10%) | 366 (9%) | 0.4483 |
| $\beta_3$ | Δ-LPS | hypermetric | 69 (15%) | 362 (9%) | **0.0002** |
| $\beta_3$ | Δ-LPS | uncertain | 370 (78%) | 3265 (80%) | 0.2016 |
| $\beta_3$ | Δ-LPS | hypometric | 36 (8%) | 433 (11%) | **0.0381** |
| $\beta_2 + \beta_3$ | LPS(+) | hypermetric | 69 (15%) | 245 (6%) | **< 0.0001** |
| $\beta_2 + \beta_3$ | LPS(+) | uncertain | 376 (79%) | 3527 (87%) | **< 0.0001** |
| $\beta_2 + \beta_3$ | LPS(+) | hypometric | 30 (6%) | 288 (7%) | 0.5701 |

### Alternate analyses demonstrate similar results for LPS-induced expression

Of the nine primate species we sampled, six had high-quality reference genomes with standardized ENSEMBL annotations, but three did not. Because it is possible that the differences in assembly and annotation quality are associated with unknown biases, we filtered out the three species with lower-quality references and conducted the same analyses as before. Results of the analyses of pooled of immune reads were consistent with the full 9-species analysis regarding LPS-induced expression: both the pool of one-to-one orthologs and the pool of full orthogroups produced 95%-credible hypermetric estimates (β_2_∼ 0.13 & 0.17). The 6-species pool of one-to-one orthologs also produced a 95%-credible hypermetric estimate for relative abundance after LPS (β_2_ + β_3_ ∼ 0.08). However, analysis of whole immune orthogroups in this reduced set of species produced a 95%-credible *hypo*metric estimate of baseline transcript abundance (β_2_ ∼ -0.13) (Table 1, Figs. S8-11). Analysis of individual gene patterns had a similar result: immune genes were significantly more likely to have hypermetric LPS-induced expression and relative abundance after LPS, but were significantly *less* likely to have hypermetric baseline relative abundances (Table S1, Data S3). These slightly different outcomes could result from greater statistical power in the smaller dataset due to the larger number of genes identified as one-to-one orthologs, or from a bias in the intercept for lower-quality reference genomes, which would add noise to baseline abundance across species but leave estimates of LPS-induced expression unbiased.

### Exaggerated LPS response in larger animals supports the Safety Factor hypothesis

Our results show that immune gene expression is indeed associated with body mass in a disproportionate (i.e., allometric) manner. Large primates increase their immune gene expression more during simulated sepsis than small primates, which may result in disproportionately greater relative abundance of immune-related gene transcripts during infection in large species (Fig. 3). This result suggests that immune defense genes have greater priority in larger animals relative to other genes, consistent with the Safety Factor hypothesis.

Genes with hypermetric responses and abundances were represented by diverse immune functions, such as pathogen and damage sensors, transcription factors, and effectors (Data S2). The allometry of immune gene transcription thus appears to be quite general. However, the mechanisms of regulation still warrant further investigation. Immune genes did vary in their individual patterns. For example, the LPS sensor toll-like receptor 4 (*TLR4*) and the pro-inflammatory cytokines interferon gamma (*IFNG*), interleukin 6 (*IL6*), and interleukin 1 beta (*IL1B*) were all estimated to have hypermetric relative abundance after LPS, consistent with the Safety Factor hypothesis and the analyses of the sum of all immune genes. The hypermetric LPS-induced expression and isometric or slightly hypometric baseline relative abundances of *TLR4*, *IFNG*, and *IL1B* were also consistent with those results (Fig. 1 D,F,H). However, *IL6* contrasted the general trend, with a strong, 95%-credible signal of hypermetric baseline relative abundance and slightly hypometric LPS-induced expression (Fig. 1G). This could be related to the complex roles that *IL6* plays in both immunity and non-immune metabolic functions, which contrasts with more strictly-immune functions of *TLR4*, *IFNG*, and *IL1B*. We also note that not all immune-annotated genes would be predicted to result in greater defense capabilities when expressed more; for example, the immune regulatory protein cytotoxic T-lymphocyte associated protein 4 (*CTLA4*) is a *negative* regulator of inflammation and immune response [29]. This gene was a clear exception to the general pattern of immune genes: it had 95%-credible hypermetric baseline relative abundance and a hypometric estimate for LPS-induced expression (Fig. 1E). The allometry of *CTLA4* was also notable because the direction of LPS response appears to reverse across the range of primate body sizes: in the smallest species, LPS exposure resulted in increased relative transcript abundance, while in the largest species, LPS exposure resulted in decreased relative transcript abundance. These patterns could have important implications for the interpretation of studies that use model organisms to draw conclusions about this gene in humans [30]. The variable patterns of these highlighted genes are non-exhaustive examples; further genes and details can be found in Data S2.

### Future directions: sex and intraspecific effects on allometry

There is extensive evidence that across most species, males tend to have weaker immune responses than females and are more vulnerable to infection [31–34]. How these conditions interact with our results, especially given the patterns of sexual dimorphism displayed by our primate study species, is unknown. Because our samples were taken only from males, it is possible that the allometric coefficients that we describe are different for females. Depending on the physical and regulatory mechanisms that drive our results, we could imagine independence between sex- and allometric-effects, or stronger allometric effects in either sex. Similarly, other intraspecific variation, such as differences in adult size or age within a species, are difficult to assess with the current data. Although we studied adult animals and included intraspecific body size effects in our linear models, these were intended primarily as controls to ensure our primary analysis was focused on the evolutionary-scale differences among species, and our design is not ideal for interpreting the corresponding coefficients [35]. Exploring these questions should be a priority in the future.

### Eco-evolutionary implications of our results

Allometry in the magnitude of immune response may affect pathogen and parasite evolution [20]. Studies have shown that host tolerance can reduce selective pressures on pathogens, allowing them to evolve increased growth rates and increased pathology upon spillover to less-tolerant hosts [36]. Evidence in favor of the Safety Factor hypothesis suggests that smaller species may indeed tend to be more tolerant of pathogens; thus, spillover from small hosts to large may tend to result in infections with more severe pathologies than spillovers in the reverse direction. However, a straight line cannot necessarily be drawn from greater immune gene expression (or greater immune cell concentrations) to better clearance of pathogens. In another recent study, we found isometry in the constitutive ability of blood to kill three very different microbes [37]. In larger animals, metabolic constraints may cause each transcript and each granulocyte to be less effective. Paired with the present results, perhaps gene expression must be engaged hypermetrically to produce functional isometry. Thus, evolutionary mechanisms may simply compensate for the physical constraints imposed by body size variation.

## Materials and Methods

### Sample collection

We collaborated with San Antonio Primate Center, the Duke Lemur Center, the Sedgwick County Zoo and the USF Health Clinic to obtain fresh whole blood samples from nine primate species, including humans, spanning more than three orders of magnitude of body mass, ranging from a 64 g *Microcebus murinus* to a 124,100 g *Pongo abelii*. Our final dataset included two aye-ayes (*Daubentonia madagascariensis*), four ring-tailed lemurs (*Lemur catta*), four tufted capuchins (*Sapajus apella*), four Rhesus macaques (*Macaca mulatta*), two olive baboons (*Papio anubis*), two hybrid olive/yellow baboons (*Papio anubis cynocephalus*, treated as *P. anubis* for all analyses), four common marmosets (*Callithrix jacchus*), eight mouse lemurs (in a single pooled sample; *M. murinus*), one human (*Homo sapiens*), and one Sumatran orangutan (*P. abelii*). We focused sampling on males to minimize sex differences in gene expression. (Fig. 2, Data S1).

A fresh, venous whole blood sample (2 mL) was collected from each individual animal by collaborators and the date and time were noted. One milliliter of fresh blood was then transferred to each of two TruCulture tubes (Myriad RBM) containing 2 mL of a proprietary media mixture (1-part blood to 2-parts media). One tube (LPS treatment) contained 1 μg/mL of 055:B5 strain LPS (782-001087) and the other (null treatment) contained only the growth media (782-001086). Each tube was then inverted three times to ensure adequate mixing and incubated at 37 °C for 8 hours. We chose 8 hours to balance the initial burst of pro-inflammatory cytokines, which peak in humans 4 hours after an LPS challenge, and the later increase of other cytokines such as IL8, which peaks at 24 hours [38]. After incubation, tubes were centrifuged at 400-500 g for 10 min. The seraplas filter was then inserted into the tube and the supernatant (plasma/media layer) was removed. The cells left at the bottom of the tube were then transferred to a new 2 mL cryovial, RNAlater (Qiagen) was added and frozen at -80°C until RNA extraction could be completed. In the case where a 2 mL blood draw was not possible (only for 1 marmoset and 1 capuchin), the ratios inside the TruCulture tube were adjusted so that the 1:2 blood:media was consistent across samples. Because of the small body mass of the mouse lemurs, we pooled blood from eight individuals and distributed that 2 mL blood pool equally across the two samples (null and LPS). De-identified blood was collected from a single human. This study was performed in accordance with Institutional Review Board Approved Protocols as well as guidelines in the 1964 Declaration of Helsinki and its later amendments, with written informed consent obtained from the patient.

### Whole blood RNA extraction

We used a Trizol-based protocol to extract RNA from primate samples using the Direct-zol RNA Microprep kit (R2060, Zymo Research). All steps were centrifuged at 13,000 g for 30 s unless otherwise noted. Briefly, 9 μl Proteinase K was added to a 300 μl sample stored in RNAlater and incubated for 30 min. Then, 2.0 mL Trizol was added to the sample and incubated while vortexing for 15 min. We then added 2.3 mL ethanol to the mixture and vortexed it. The sample was added to a Zymo-spin IC Column fitted with a collection tube in 700 μl increments (the maximum loading volume of the column). After each increment, the sample was centrifuged, and the flow-through discarded. We then performed the DNase treatment steps. We added 400 μl RNA wash buffer and centrifuged the sample, and then added 5 μl DNAse 1 and 35 μl DNA digestion buffer and incubated it for 15 min. We added 400 μl Direct-zol RNA prewash buffer to the column, centrifuged and discarded flow-through (x2). Then 700 μl of RNA wash buffer was added to the column and centrifuged for 1.5 min to ensure removal of buffers. Finally, 10 μl of RNase-free water was then added directly to the column matrix, incubated for 1 min, and centrifuged. Concentrations and quality of each sample were checked via RNA broad range Qubit fluorometric quantification (Q10210, Thermo Fisher Scientific) and Agilent tape station 2200 (Agilent), respectively.

### Library preparation and RNAseq

Extracted RNA samples were submitted to the University of Illinois at Urbana-Champaign Roy J. Carver Biotechnology Center for library preparation. Libraries were prepared with the Ovation SOLO RNAseq kit (M01406, Tecan) with probes for human rRNA and globin RNA. They were sequenced with the NovaSeq 6000 with an S4 flow-cell and 150 bp paired-end reads. Reads were quality-trimmed before delivery and further analysis, resulting in an average of 90 bp.

### Reference transcriptomes, orthogroups, and gene expression quantification

ENSEMBL (release 109) [39] reference transcriptomes (*.cdna.all.fa.gz) were used for *M. murinus*, *C. jacchus*, *P. anubis*, *M. mulatta*, *P. abelii*, and *H. sapiens*. Because *L. catta*, *D*. *madagascariensis*, and *S*. *apella* are not yet integrated into ENSEMBL, reference transcriptomes were created with the following workflow. First, NCBI GenBank genomes were acquired (GCF_020740605.2, GCA_023783475.1, and GCF_009761245.1, respectively), and HISAT2 v2.1.0 [40] indexes were built. Then, sequencing reads for each sample were assembled into ‘superreads’ using the superreads.pl script provided for use with StringTie v1 [41]. Superreads and unassembled paired-end reads were then mapped to their respective genome using HISAT2 with options --no-discordant --no-mixed --dta-cufflinks. StringTie v2.2.1[42] was then used to generate transcriptomes with default settings. TransDecoder v5.5.0 [43] was used to predict peptide sequences. The longest predicted peptide was isolated for each gene, and these peptides were input to Orthofinder v2.5.4 [44] in combination with ENSEMBL reference peptides (filtered from Compara.109.protein_default.aa.fasta.gz) to identify orthogroups across all nine species. The phylogeny from Hedges *et al.* 2015 was subsetted and used as a fixed species tree in the Orthofinder analysis. Orthogroups were identified using Orthofinder’s root-level ‘Phylogenetic_Hierarchical_Orthogroups’ output file (N0.tsv), and and 1:1 orthologs were determined by filtering the rows of this table. Gene expression estimates were generated using Salmon v1.6.0 [45] with default settings and raw sequencing reads from each sample.

### Differential abundance analysis of genes

To assess differential abundance of each gene, count data were analyzed with phylogenetic generalized linear mixed models using the function pglmm() from the R package *phyr* v1.1.0 [46]. To approximately account for the discrete and compositional nature of RNASeq count data, a Poisson likelihood with a log link was used with an offset term unique to each observation. This offset was determined by importing Salmon quantification outputs into an R object with the package *tximport* v1.26.1 [47], filtering the genes to include only 1:1 orthologs, and using the DESeq2 v1.38.2 [48] function *estimateSizeFactors*. The resulting ‘normalization factors’ are designed to compensate for varying sequencing depth, compositionality, and average transcript length across samples. Binary LPS treatment, the standardized log of each species’ average body mass [28], the standardized log of species-normalized individual body size, and the interaction of LPS treatment with each body size effect, were modeled as fixed effects (β_1_-β_5_), with random effects for individual, independent and identically distributed (i.i.d.) species, species phylogeny, and the interactions between the treatment and i.i.d. species and species phylogeny. *phyr* also automatically includes an observation-level random effect to account for overdispersion relative to the expected variance of the Poisson likelihood. The aforementioned phylogenetic tree from Hedges *et al.* 2015 [27] was used for phylogenetic random effects. In this model, the main effect of LPS treatment (β_1_) can be interpreted as the change in expression after LPS treatment for each gene at the ancestral body size (somewhat arbitrary for the purposes of this study, and determined by optimization with the *ace* function from the R package *ape* 5.7-1 [49]), the main effect of species’ average body mass (β_2_) can be interpreted as the allometric effect on the baseline relative abundance of each gene (e.g., LPS(-)), the interaction between LPS treatment and species’ average body mass (β_3_) can be interpreted as the allometric effect on change in expression after LPS treatment (e.g., Δ-LPS), the main effect of species-normalized individual body size (β_4_) may be caused by either aging-related changes or within-species allometric differences in the baseline relative abundance of each gene, and the interaction between LPS treatment and species-normalized individual body size (β_5_) may be caused by either aging-related changes or within-species allometric differences in the change in expression after LPS treatment. The random effect for individuals enables the model to focus on paired differences between and LPS and null treatment, the main species effects control for incidentally correlated evolution of body mass and baseline gene expression, and the species interaction effects control for incidentally correlated evolution of body mass and the change in expression during LPS treatment. The model was fit using approximate Bayesian inference with default priors with *INLA* v21.02.23 [50]. The sums of β_2_ and β_3_ and of β_4_ and β_5_ were calculated during model fitting using *INLA*’s ‘lincomb’ option, to represent the actual allometric slopes among LPS-treated samples (e.g., LPS(+)). Models that did not converge were retried up to 15 times, and genes were filtered from further analysis if they did not converge. These genes (55 out of 4590 1:1 orthologs) tended to have some samples with low counts, which can exacerbate overfitting and lead to singular model matrices. Fixed effect coefficients were extracted, and individual genes were determined to have significant differential relative abundance if their 95% credible interval did not contain zero. Two-by-two contingency tables were constructed summarizing the number of genes that were allometric (or not) and immune-annotated (or not), and Fisher’s exact tests were used to determine if each allometric pattern was more likely in immune vs. non-immune genes.

### Differential abundance analysis of total immune genes

Orthologous gene sets were annotated with their corresponding immune functions as defined by Deschamps *et al*. 2016 [51]. External gene names were determined for each ortholog by querying the Ensembl BioMart [39] with the R package *biomaRt* v2.54.1 [52], using human members of each orthogroup, and these gene names were used to extract immune function annotations from Table S1 from Deschamps *et al*. Two workflows were used to assess the total expression of immune genes. In the first, counts for all immune-annotated 1:1 orthologs were summed, and average gene lengths for the pool were calculated for use in DESeq2’s normalization factor calculations. This method is conservative in focusing only on identifiable 1:1 orthologs and can utilize observation-specific normalization factors so that results may be interpreted as differences in the number of transcripts across samples. In the second, counts of all genes that belonged to immune-annotated orthogroups were pooled, and ‘size factors’ that were consistent for a given sample were calculated by taking the geometric mean of the normalization factors of all 1:1 orthologs. This method is more comprehensive, covering all genes with likely immune functions, but because it does not compensate for differences in average transcript length across samples, it is *less confidently* interpretable as differences in numbers of transcripts. The choice to use ‘size factors’ instead of ‘normalization factors’ here was made because it is likely that gene predictions are fragmented for the three species with lower-quality references, and apparent gene duplications or losses in these species could artificially influence the predicted average transcript length. This version is better interpreted as the amount of RNA with a given annotation, rather than the abundance of transcripts with a given annotation. For both analyses, total immune gene abundance was modeled with *phyr* and *INLA* using the same protocol and formula as analyses of individual genes. Model diagnostics were performed using *DHARMa* [53]

To analyze only the six species with higher-quality ENSEMBL reference genomes, all differential abundance methods were repeated after filtering the previous Salmon and Orthofinder outputs.

All analyses that used R throughout the manuscript used version 4.2.2 [54]. The packages *ggtree* v3.6.2 [55] and *ggplot2* v3.4.1 [56] were used for plotting.

## Acknowledgements

The authors thank Debbie Christian, Deborah Chavez, and Chris Chen at the Southwest National Primate Research Center at the Texas Biomedical Research Institute; Erin Ehmke, Kay Welser, and Megan Davison at the Duke Lemur Center; and Jennifer Kropf and Sandy Wilson at the Sedgwick County Zoo for their collaboration and support in obtaining and processing these primate samples. We thank Manfred Schmolz at Myriad RBM for help with TruCulture techniques and Alvaro Hernandez in troubleshooting RNA extractions and sequencing. Finally, thanks to Samantha Oakey, Matthew Mercurio, Min Zhang, and the University of South Florida Genomics Program Sequencing Core and Computational Core/Omics Hub for laboratory and computational support.

## Author contributions

Conceptualization: CJD, LBM, RHYJ, RM

Data curation: RM, ECR, SRA, RAM

Formal analysis: RM

Funding acquisition: CJD, RHYJ, LBM

Investigation: ECR, SRA

Methodology: RM, ECR, SRA, CJD, LBM, RHYJ

Project administration: CJD, LBM

Resources: ECR, CJD, LBM, JWL

Supervision: CJD, LBM, RHYJ Visualization: RM

Writing – original draft: RM, CJD, LBM

Writing – review & editing: RM, CJD, LBM, RHYJ, RAM, ECR, SRA, JWL

## Data and materials availability

Quality-trimmed RNASeq data is available in NCBI BioProject PRJNA834816. All code used for analysis is available at https://github.com/rmcminds/McMinds_et_al_2022 and at Zenodo (doi: 10.5281/zenodo.10779069).

## Supplementary Materials

**Fig. S1.**
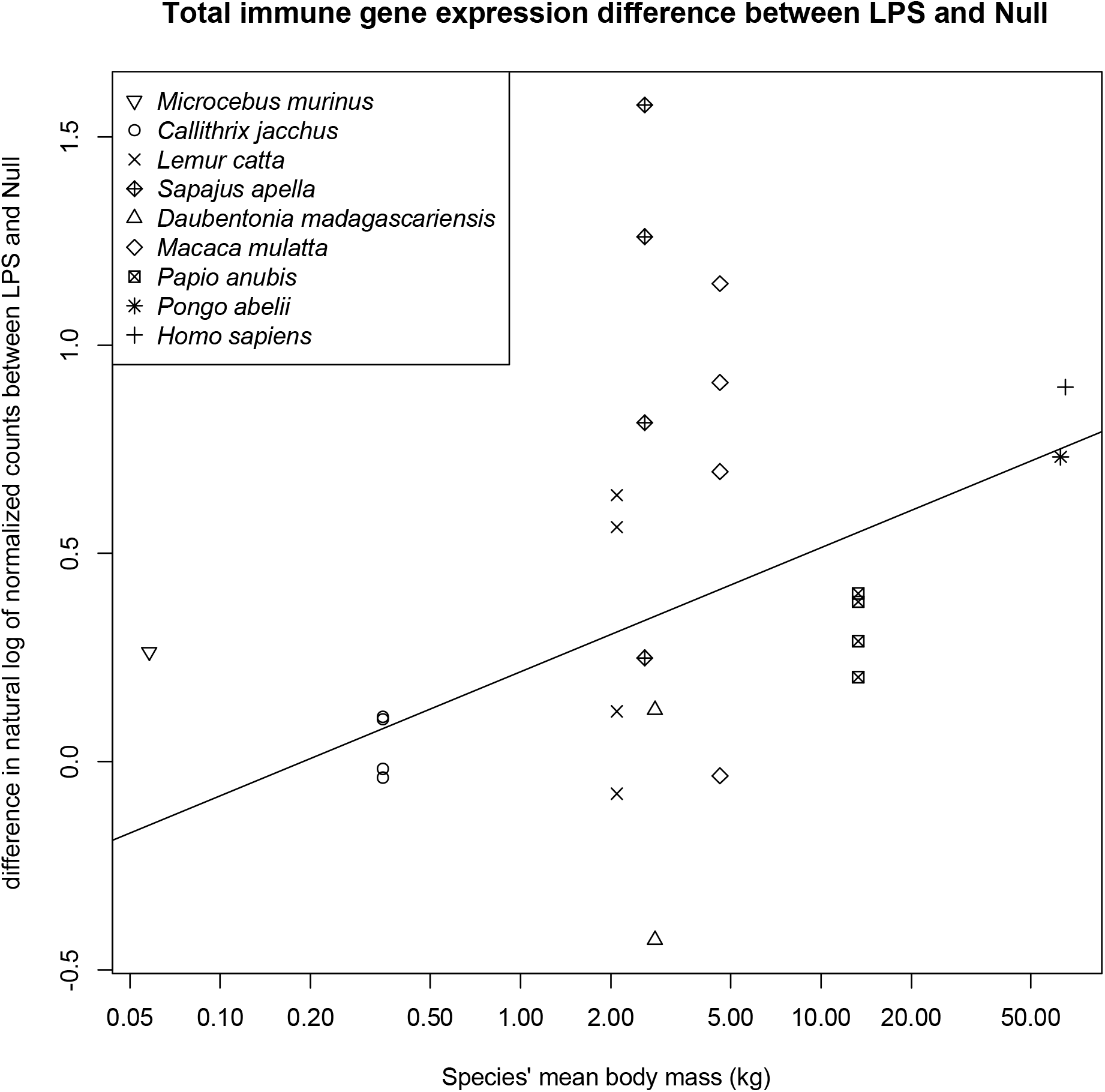
Summed counts of all immune-annotated 1:1 orthologous genes show hypermetric response to sepsis. This figure draws on the same data as Figure 3 in the main text, but shows the difference between LPS- and Null-treated samples from each individual rather than the raw abundance values. Body mass is plotted on the log scale.

**Fig S2.**
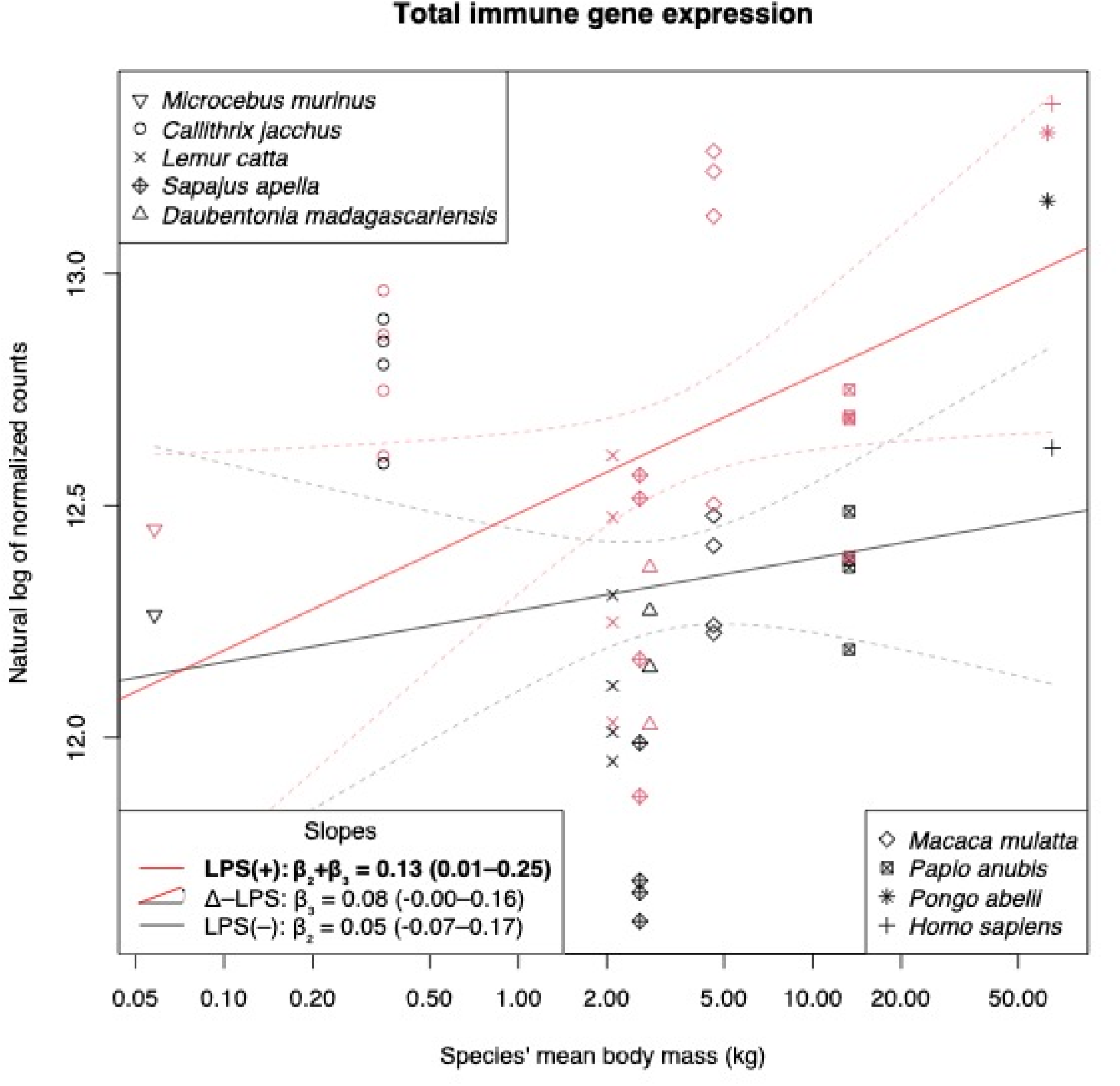
Summed counts of all immune-annotated reads show hypermetric concentrations after LPS. This figure is analogous to Figure 3 in the main text, but is the sum of all reads annotated with immune functions, even if they belong to orthogroups that had duplications or gene deletions. This analysis is more comprehensive as it was able to include more genes (>4000 compared to >1200), but it is probably more susceptible to differences in gene lengths, or apparent gene lengths due to lower quality transcriptome assemblies. As an alternate method with different assumptions that shows something similar, it’s a nice backup. Theoretically, this figure might be best interpreted as an increase in total immune-related RNA, whereas Fig. 3 depicts an increase in the total number of immune-annotated transcripts (but in the absence of widescale biases in gene or transcript lengths, the interpretation should be similar).

**Fig. S3.**
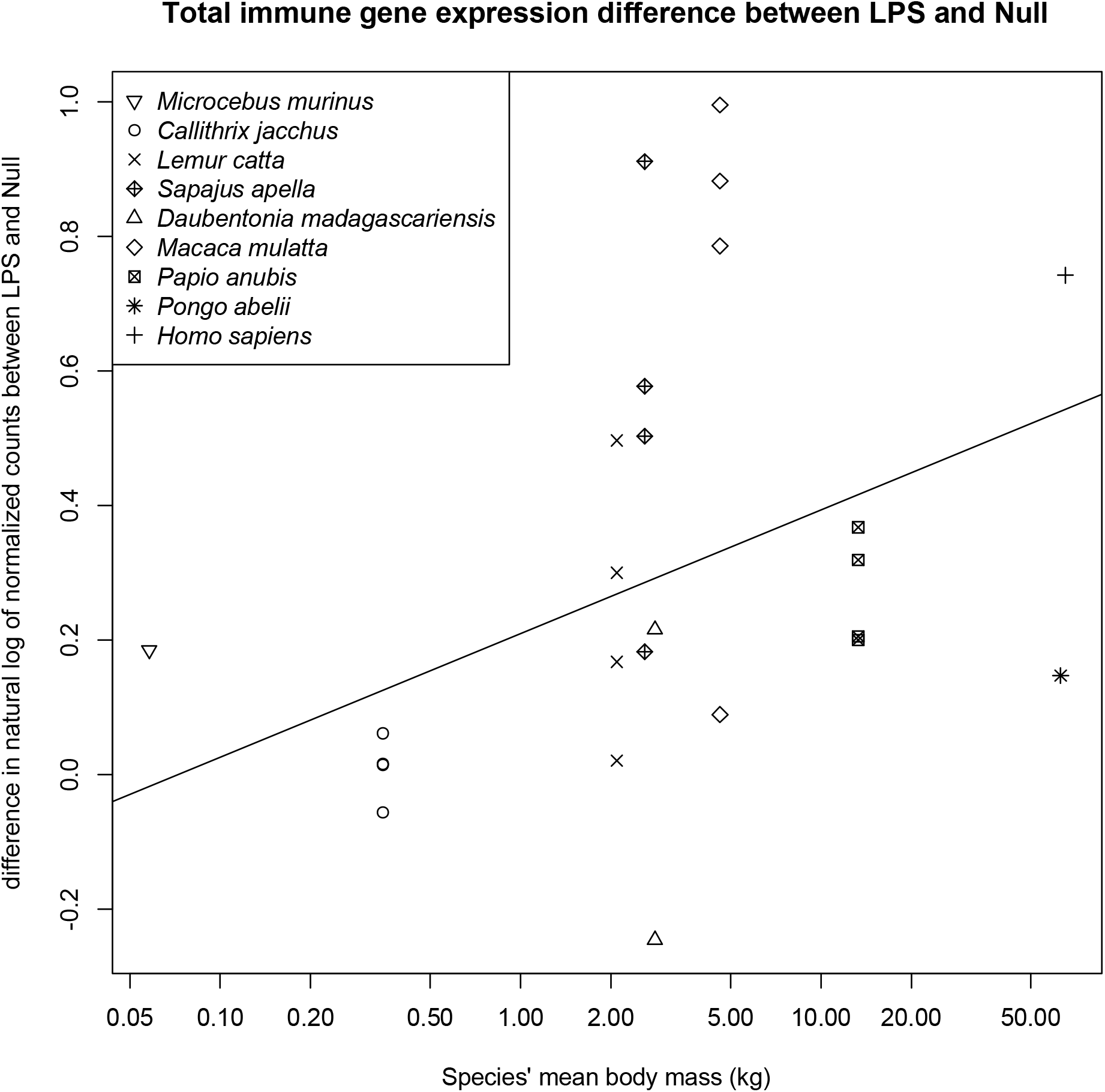
Sum of all counts of reads that belong to immune-annotated orthogroups. This figure is the same as Figure S1, but this figure includes all immune reads rather than just 1:1 orthologs.

**Fig. S4.**
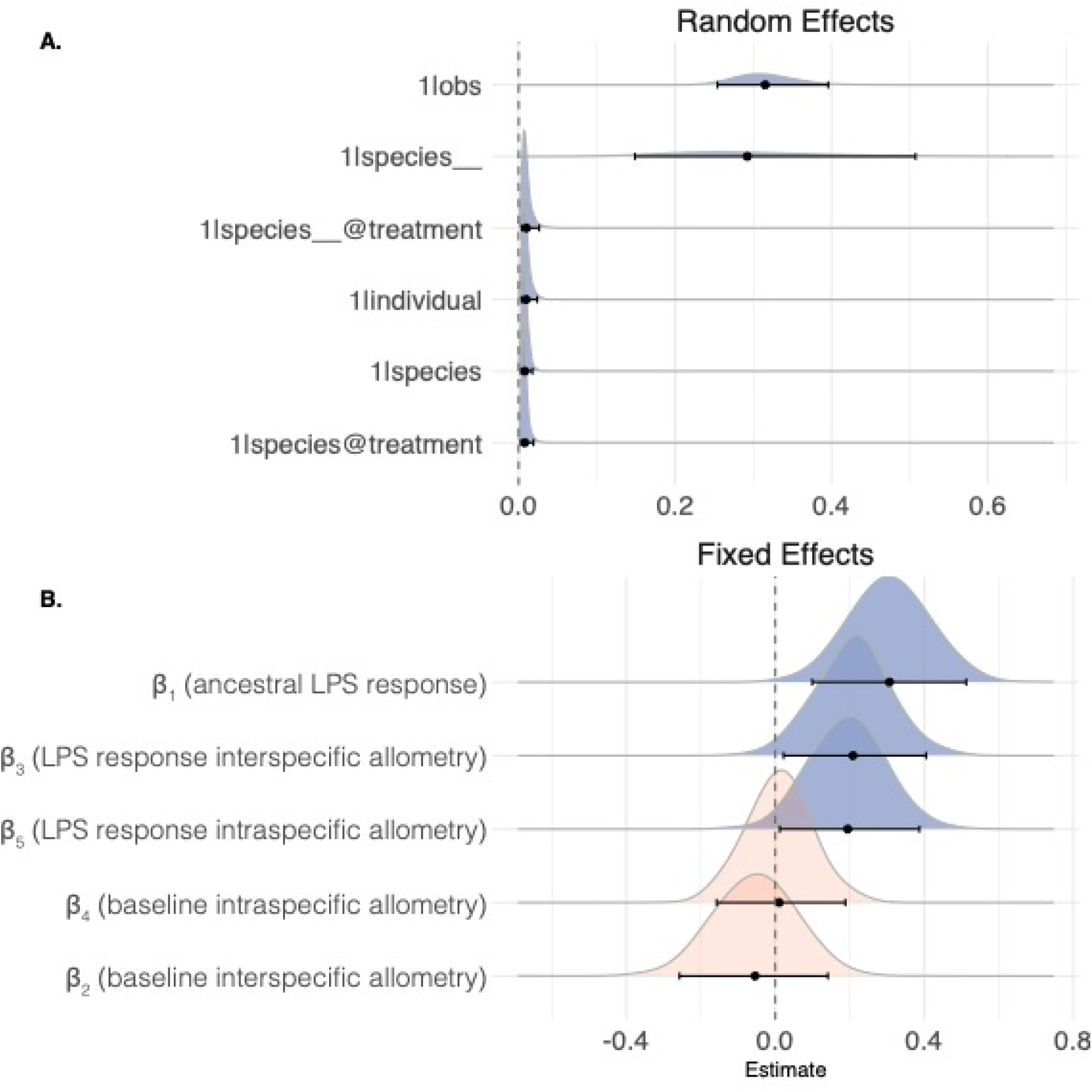
Density plots for 9-species 1:1 ortholog analysis. **(A)** Posterior distributions of the residual variance for each random effect. Effects are ordered by median estimates. Names are in *phyr* notation: 1|obs is the observation-level extra-poisson variance; 1|species is the (Brownian) phylogenetic effect on baseline abundance; 1|species @treatment is the phylogenetic effect on LPS-induced expression; 1|individual is the effect of individual; 1|species is the non-phylogenetic effect of species on baseline abundance; and 1|species@treatment is the non-phylogenetic effect of species on LPS-induced expression. **(B)** Posterior distributions of the fixed effect coefficients. The grand intercept (alpha) has been removed, and coefficients are ordered by median estimates. The median and 95% credible intervals are shown on each axis. Density plots are blue if the 95% credible interval does not include zero.

**Fig. S5.**
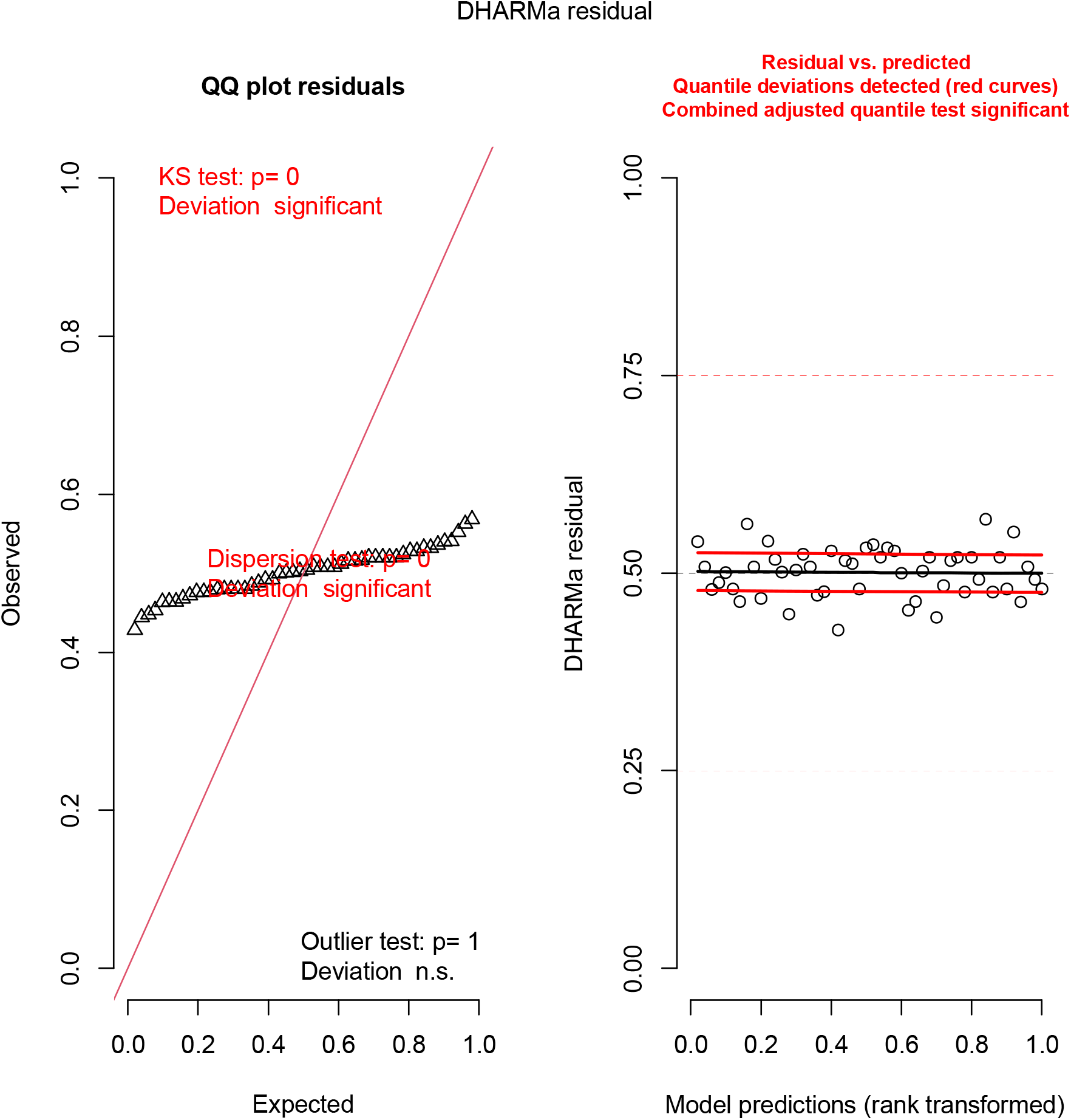
DHARMa plots for 9-species 1:1 ortholog analysis. The significant results from the Kolgorov-Smirnov and dispersion tests are consistent with an overfit model applied to count data. This outcome results from the ability of the continuous parameters, including observation-level random effects, to predict each observation almost perfectly, leaving little residual variance to be explained by the Poisson count model. This result is common for Poisson models with observation-level random effects and is usually associated with lower statistical power. This suggests that our analysis is conservative and should not strongly influence the interpretation of effects that are found to be significant.

**Fig. S6.**
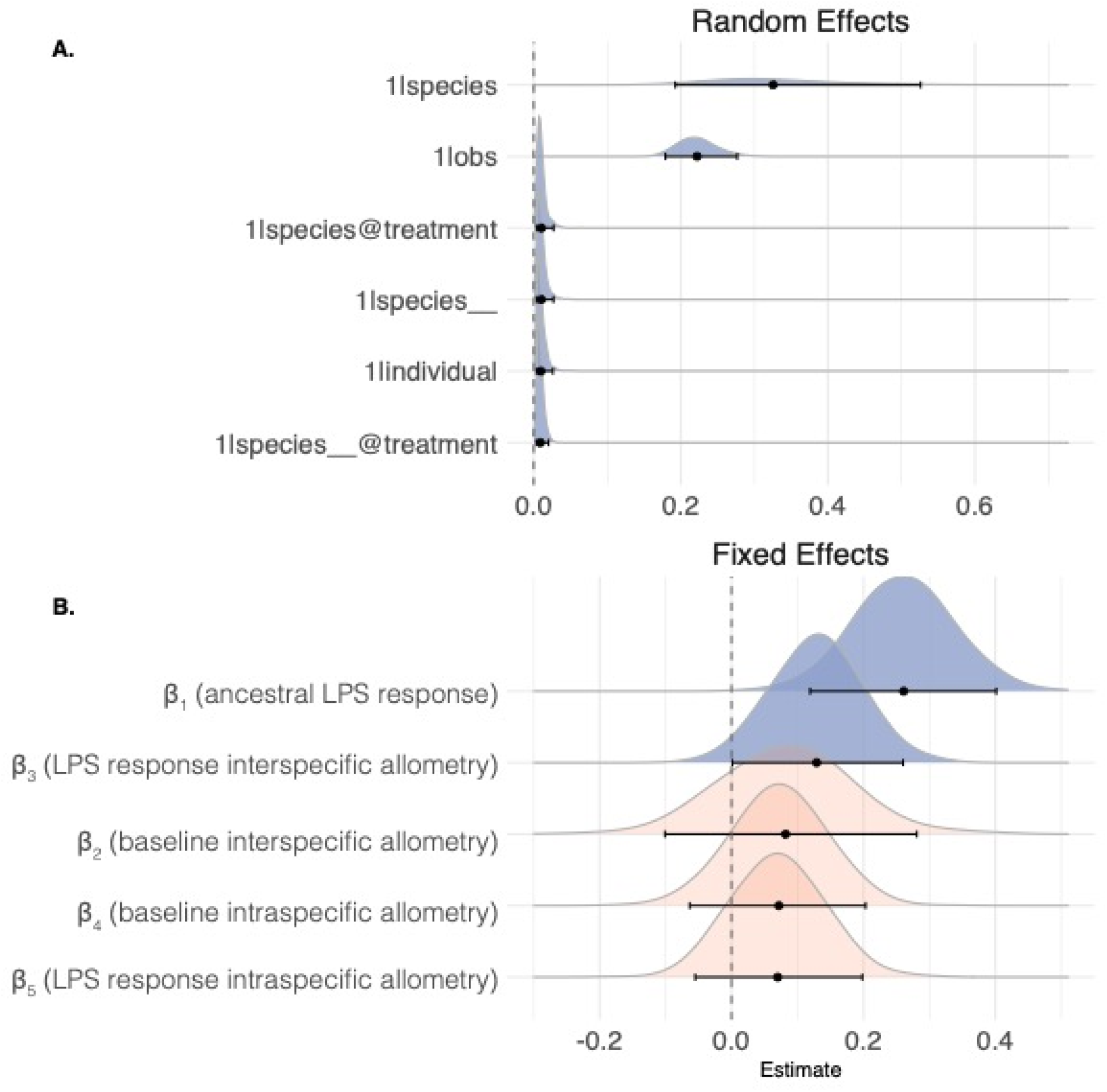
Density plots for 9-species orthogroup analysis. **(A)** Posterior distributions of the residual variance for each random effect. Effects are ordered by median estimates. Names are in *phyr* notation: 1|obs is the observation-level extra-poisson variance; 1|species is the (Brownian) phylogenetic effect on baseline abundance; 1|species @treatment is the phylogenetic effect on LPS-induced expression; 1|individual is the effect of individual; 1|species is the non-phylogenetic effect of species on baseline abundance; and 1|species@treatment is the non-phylogenetic effect of species on LPS-induced expression. **(B)** Posterior distributions of the fixed effect coefficients. The grand intercept (alpha) has been removed, and coefficients are ordered by median estimates. The median and 95% credible intervals are shown on each axis. Density plots are blue if the 95% credible interval does not include zero.

**Fig. S7.**
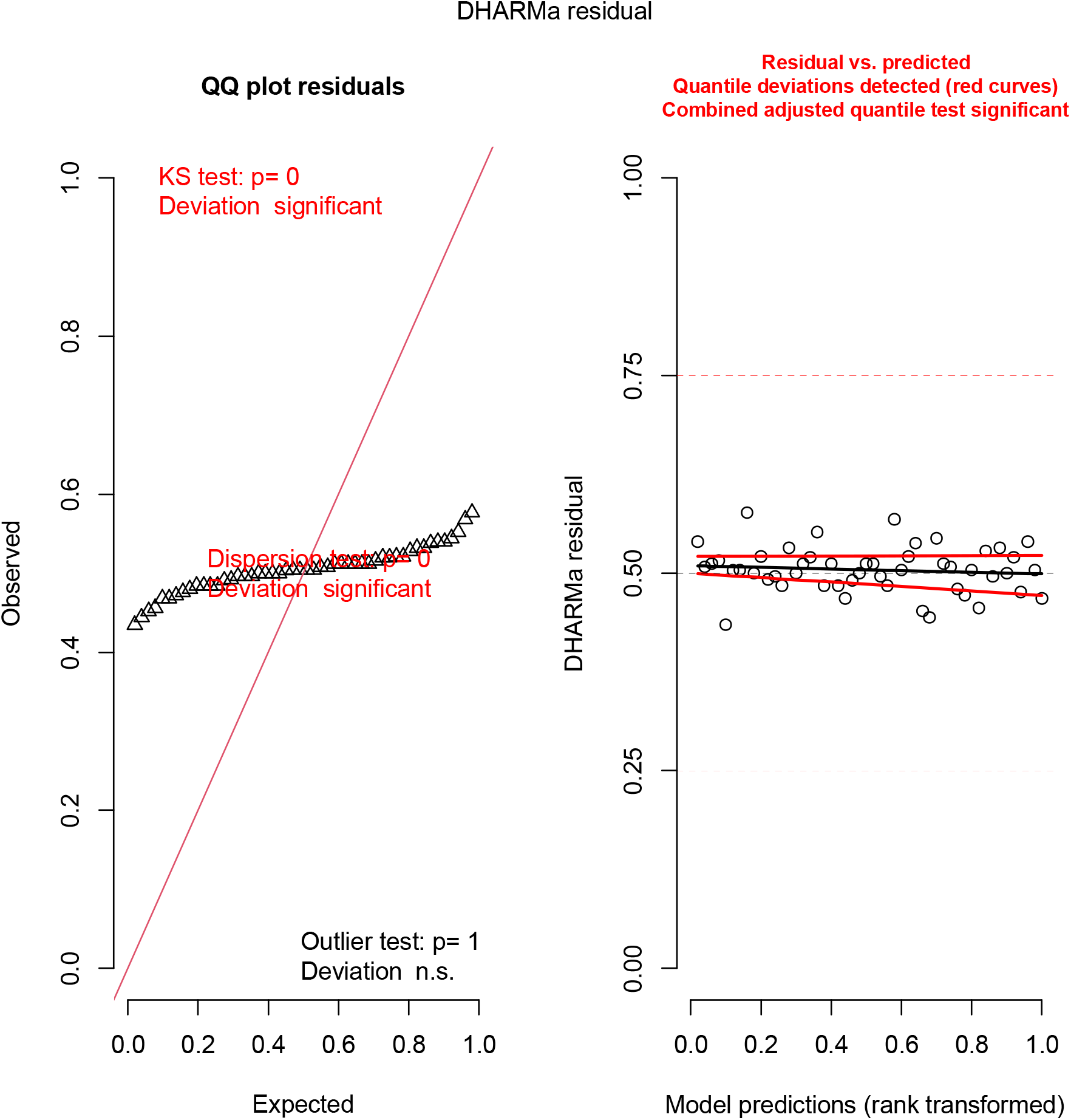
DHARMa plots for 9-species orthogroup analysis. This is analogous to Fig S5, with similar interpretations of overfitting and the conservative nature of the tests.

**Table S1.** Number of immune- or non-immune-annotated genes with hypometric, hypermetric, or uncertain relation to body size (6-species analysis). Genes were considered hypometric in each context (baseline abundance = LPS(-), LPS-induced expression = Δ-LPS, and abundance after LPS = LPS(+)) if the pGLMM 95% credible interval for the corresponding beta was entirely negative, hypermetric if it was entirely positive, and uncertain if it included zero. Innate immune annotations were derived from Deschamps *et al*. 2016. P-values correspond to Fisher’s exact test comparing the proportions of immune or non-immune genes with each allometric pattern vs. the sum of the other two patterns.

| Parameter | Gene expression context | Allometry | Immune genes | Non-immune genes | P-value (Fisher) |
| --- | --- | --- | --- | --- | --- |
| $\beta_2$ | LPS(-) | hypermetric | 186 (19%) | 2494 (24%) | **0.0012** |
| $\beta_2$ | LPS(-) | uncertain | 674 (69%) | 7010 (66%) | 0.0824 |
| $\beta_2$ | LPS(-) | hypometric | 116 (12%) | 1071 (10%) | 0.0876 |
| $\beta_3$ | Δ-LPS | hypermetric | 116 (12%) | 765 (7%) | **< 0.0001** |
| $\beta_3$ | Δ-LPS | uncertain | 751 (77%) | 8136 (77%) | 1.0000 |
| $\beta_3$ | Δ-LPS | hypometric | 109 (11%) | 1674 (16%) | **0.0001** |
| $\beta_2 + \beta_3$ | LPS(+) | hypermetric | 129 (15%) | 676 (6%) | **< 0.0001** |
| $\beta_2 + \beta_3$ | LPS(+) | uncertain | 770 (79%) | 8948 (85%) | **< 0.0001** |
| $\beta_2 + \beta_3$ | LPS(+) | hypometric | 77 (6%) | 951 (9%) | 0.2644 |

**Fig. S8.**
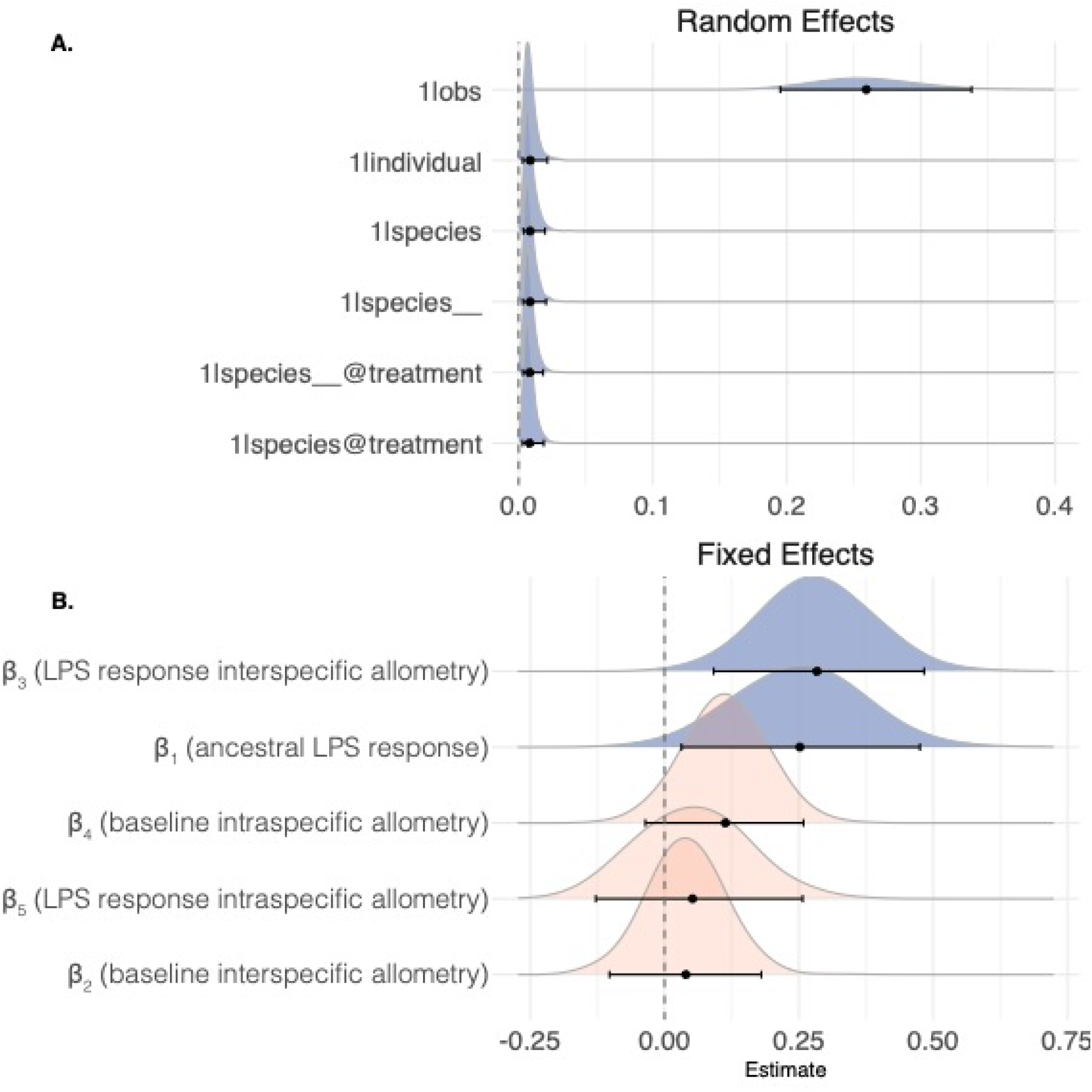
Density plots for 6-species 1:1 ortholog analysis. **(A)** Posterior distributions of the residual variance for each random effect. Effects are ordered by median estimates. Names are in *phyr* notation: 1|obs is the observation-level extra-poisson variance; 1|species is the (Brownian) phylogenetic effect on baseline abundance; 1|species @treatment is the phylogenetic effect on LPS-induced expression; 1|individual is the effect of individual; 1|species is the non-phylogenetic effect of species on baseline abundance; and 1|species@treatment is the non-phylogenetic effect of species on LPS-induced expression. **(B)** Posterior distributions of the fixed effect coefficients. The grand intercept (alpha) has been removed, and coefficients are ordered by median estimates. The median and 95% credible intervals are shown on each axis. Density plots are blue if the 95% credible interval does not include zero.

**Fig. S9.**
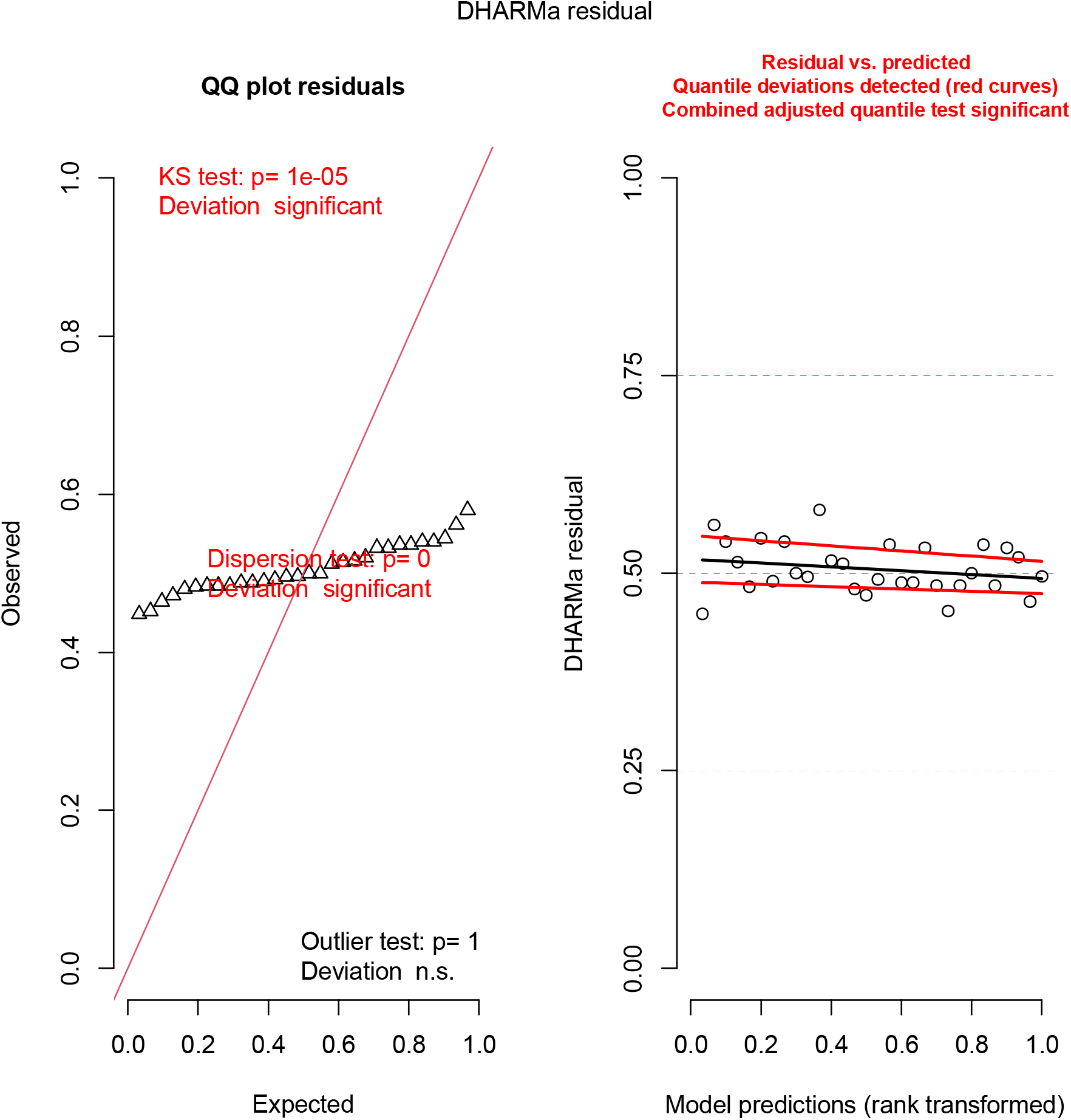
DHARMa plots for 6-species 1:1 ortholog analysis. This is analogous to Fig S5, with similar interpretations of overfitting and the conservative nature of the tests.

**Fig. S10.**
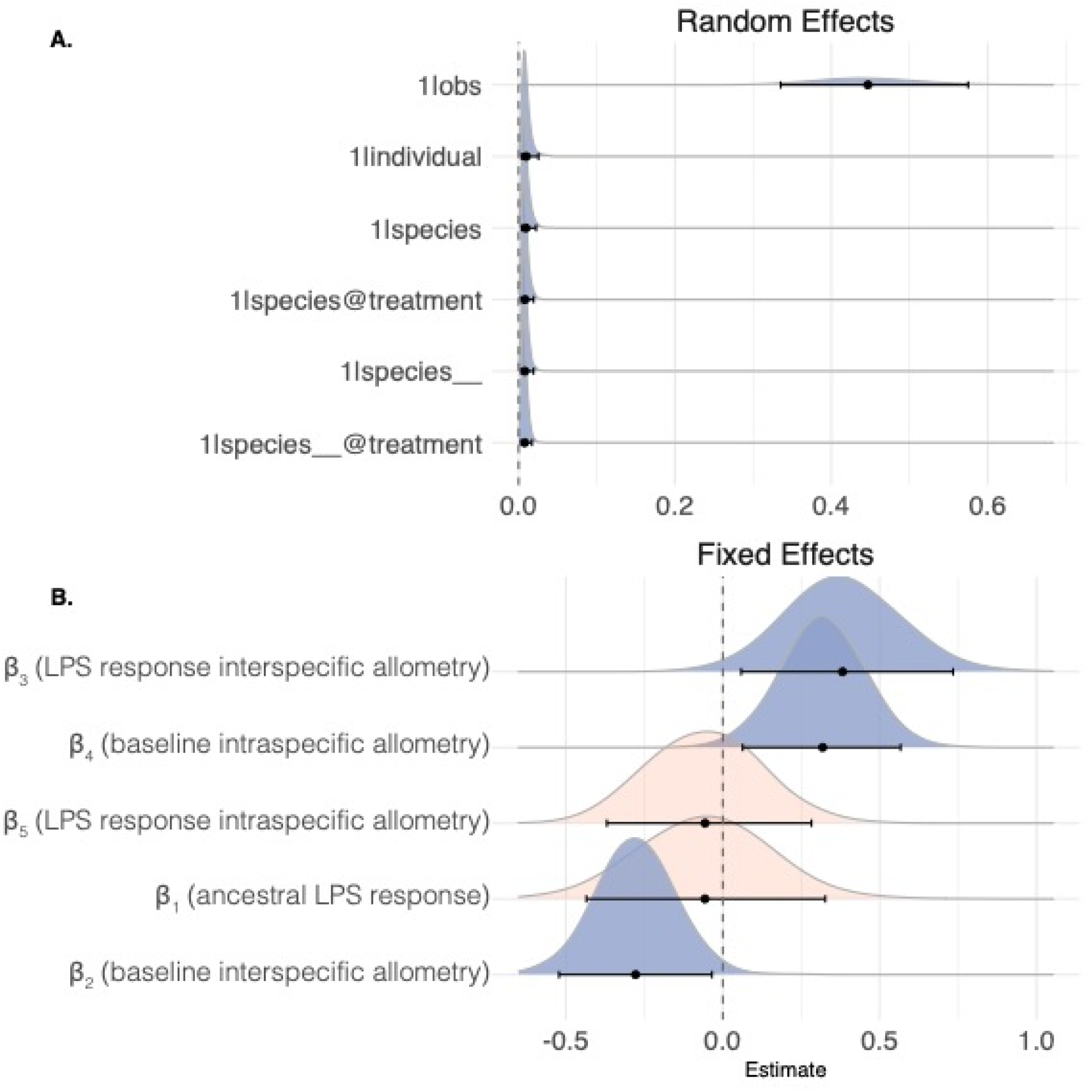
Density plots for 6-species orthogroup analysis. **(A)** Posterior distributions of the residual variance for each random effect. Effects are ordered by median estimates. Names are in *phyr* notation: 1|obs is the observation-level extra-poisson variance; 1|species is the (Brownian) phylogenetic effect on baseline abundance; 1|species @treatment is the phylogenetic effect on LPS-induced expression; 1|individual is the effect of individual; 1|species is the non-phylogenetic effect of species on baseline abundance; and 1|species@treatment is the non-phylogenetic effect of species on LPS-induced expression. **(B)** Posterior distributions of the fixed effect coefficients. The grand intercept (alpha) has been removed, and coefficients are ordered by median estimates. The median and 95% credible intervals are shown on each axis. Density plots are blue if the 95% credible interval does not include zero.

**Fig. S11.**
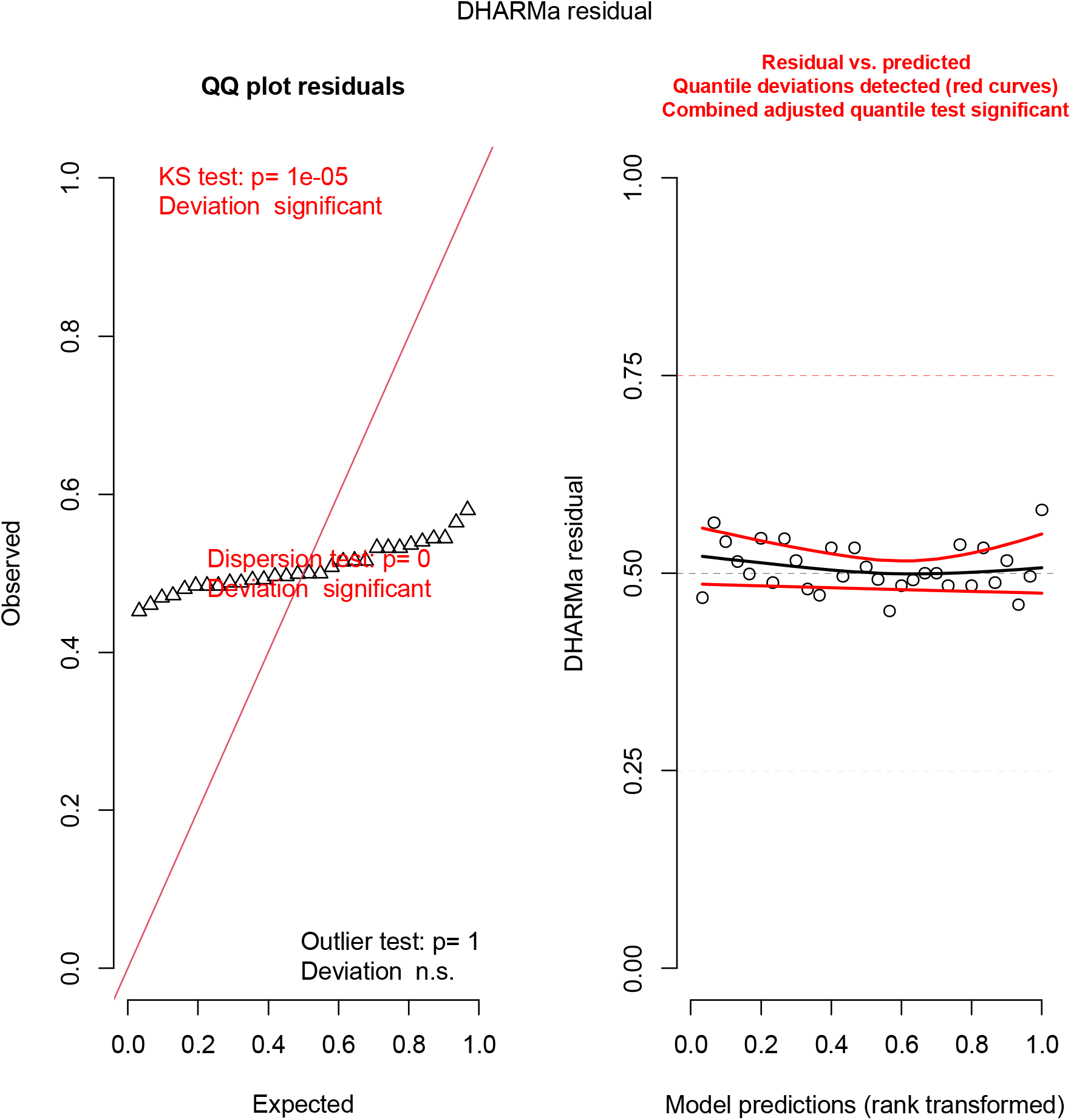
DHARMa plots for 6-species orthogroup analysis. This is analogous to Fig S5, with similar interpretations of overfitting and the conservative nature of the tests.

Data S1 to S3

